# Evolving Insights from *SARS-CoV-2* Genome from 200K COVID-19 Patients

**DOI:** 10.1101/2021.01.21.427574

**Authors:** Sanket Desai, Aishwarya Rane, Asim Joshi, Amit Dutt

## Abstract

We present an updated version of our automated computational pipeline, Infection Pathogen Detector IPD 2.0 with a *SARS-CoV-2* module, to perform genomic analysis to understand the pathogenesis and virulence of the virus. Analysing the currently available 208911 SARS-CoV2 genome sequences (as accessed on 28 Dec 2020), we generate an extensive database of sample-wise variants and clade annotation, which forms the core of the *SARS-CoV-2* analysis module of the analysis pipeline. A comparative account of lineage-specific mutations in the newer *SARS-CoV-2* strains emerging in the UK, South Africa and Brazil along with data reported from India identify overlapping and lineages specific acquired mutations suggesting a repetitive convergent and adaptive evolution. Thus, the persistence of pandemic may lead to the emergence of newer regional strains with improved fitness. IPD 2.0 also adopts the recent dynamic clade nomenclature and shows improvement in accuracy of clade assignment, processing time and portability, to its predecessor and thus could be a vital tool to help facilitate genomic surveillance in a population to identify variants involved in breakthrough infections.

## Introduction

The *SARS-CoV-2* is mutating and evolving with time and geographical distribution, as typical of any RNA virus, indicating generation of an increasing pool of emerging diversity in the viral strains (1). The mutation accrual may indicate natural purifying selection adding to the fitness of virus that may impact changes in the outcome of COVID-19 disease and its transmissibility (2). The emergence of newer variants with higher infectivity or potential to impact vaccine efficacy underlines the significance of enhancing efforts to sequence the genome of the virus from across the globe.

Genome sequencing of *SARS-CoV-2* is the most widely used method for tracking strains and identification of novel emerging variants in the population. With the generation of an increasing pool of emerging diversity in the viral strains with time and geographical distribution, several national initiatives such as the *SARS-CoV-2* Sequencing for Public Health Emergency Response, Epidemiology and Surveillance (SPHERES) (3), COVID-19 Genomics UK Consortium (COG-UK) (4) and Indian *SARS-CoV-2* Genomics Consortium (INSACOG) (5) have enacted dynamic genomic surveillance to identify novel region-specific variants involved in breakthrough infections (6). Even a modest increase in infectivity rate of a regional variant or a reduction in vaccine efficacy or increased transmission would require immediate stringent measures to be put in place to contain the spread of the strain. Thus, automated measures are needed to perform integrated analysis to identify the newer variants.

We recently developed a computational tool, Infectious Pathogen Detector (IPD) with a *SARS-CoV-2* module to determine the abundance, mutation rate and phylogeny of the *SARS-CoV-2* genome, from the heterogeneous advanced sequencing data (7). With the evolving nomenclature of the *SARS-CoV-2* clades (8) and an exponential increment in the *SARS-CoV-2* genomic variant data, we aimed to expand the variant database and revise the clade assessment module for IPD 2.0. Our variant analysis, performed to create an updated *SARS-CoV-2* variation database, reveal a uniform distribution of variants across the genome, with selective enrichment of variants at hotspot regions. Additionally, we extended our analysis to include the emerging strains, B1.1.7, B1.135 and P1, and present a comparative account of recurrent mutations among these strains against the Indian variant pool, to determine any pre-existing variants from the novel strains. From the generated database, using IPD 2.0, we further evaluate the clade assessment accuracy and improvement in run-time to its predecessor.

## Materials and methods

Complete, high coverage *SARS-CoV-2* genome sequences with length greater than 29000 bp and their metadata were downloaded from GISAID database (9). In total 208911 *SARS-CoV-2* genome sequences were downloaded (as of December 28, 2020). The genome sequences having ‘N’ at the ends were trimmed and once having a length greater than 29,000 bp were retained. The filtered genome sequences (n=200865) were used for further analysis. Variant calling was performed using Snippy (10), on the filtered/cleaned sequences (n=200865), using the Wuhan strain sequence (Genbank ID NC_045512) as the reference genome. The tabular annotated variant output files for individual samples, obtained from Snippy were combined using custom scripts to retain sample, EPI ID and date information into the tabular file. Post-processing, statistical evaluation and gene-wise analysis of the data was performed in the R programming environment. The *SARS-CoV-2* genome sequences (n=200865) were subject to clade analysis using NextClade program in the NextStrain toolkit (11). Sequence data for variant analysis of new emerging strains, UK (B 1.1.7) and South African (B 1.315) sample sequences, were separately downloaded from GISAID, whereas the sequences for Brazil strain (P.1) were obtained from (12). Variant calling was performed using Snippy and the obtained variants were merged to generate a list of private variants represented in at least 50 percent of the sequences for a particular strain. This list was further used for comparison with Indian variant dataset.

The annotated, tabular, sample merged files generated upon post-processing of Snippy output were subject to mutation profile analysis. Mutation profiles consisting of a unique set of mutations in individual samples were generated. Using a representative sample / GISAID ID for individual profile the variants were extracted from the tabular variant data. The analysis generated 99,301 representative mutation profiles. The tabular data for variants (with representative sample information) was indexed using Tabix (13) and this indexed database is integrated into IPD 2.0 to identify the novel variants in the analysed samples. The clade and sub-clade variant set provided by the NextStrain, are also a part of the IPD 2.0 clade assessment program. The algorithm for novel variant identification, variant profile comparison and clade assignment is described earlier (7). We further tested the accuracy of the clade assessment module of IPD 2.0 using the simulated dataset from the 8 representative clades (described in Supplementary methods).

In order to further ease the process of installation of IPD on the host machine, we have changed the build and installation process for IPD 2.0, wherein we provide a conda based environment which is installable across all the Unix platforms with Miniconda. We also provide the pre-compiled compressed reference data (including the variant database) required to run IPD 2.0 (refer user manual on http://ipd.actrec.gov.in/ipdweb/manual.html). This allows users to bypass the process of installation of dependencies on the host machine and makes IPD 2.0 portable across variants of Unix platforms.

## Results

We analyzed the *SARS-CoV-2* genome data to generate a comprehensive variant dataset for *SARS-CoV-2* genome. We examined 2.58 million *SARS-CoV-2* mutations found in 200865 samples from 155 different countries, in the sequences downloaded from GISAID (9) (as accessed on 28 Dec 2020). In this dataset, in comparison to the ancestral reference *SARS-CoV-2* Wuhan strain (14), we find 1004453 (38.88%) synonymous, 1327548 (51.39 %) nonsynonymous mutations and 242631 (9.39%) mutations in the intergenic region comprising of coding 5’ and 3’UTRs, indicating a relatively higher representation of nonsynonymous mutations. Among nonsynonymous mutations, missense mutations (49.54%) were more frequent than stop lost (1.17%), stop gain (0.66%) and deletions/ insertions (0.23%). In overall, 6.6 nonsynonymous, 5 synonymous and 1.20 intergenic mutations per sample were observed, as shown in Table 1.

**Table 1:**
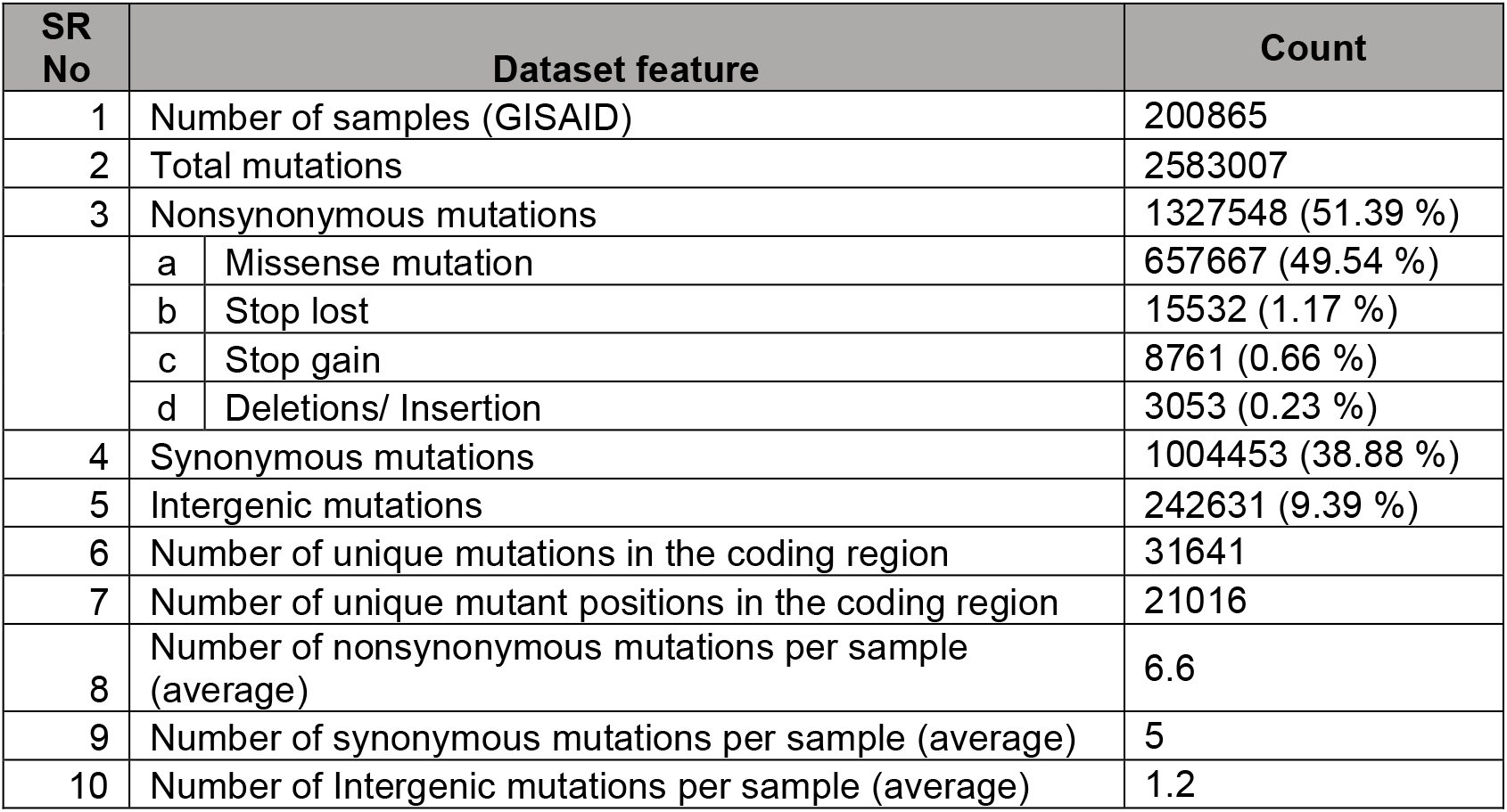
Distribution of type of *SARS-CoV-2* genome variants across the sequences in GISAID database

| SR No | Dataset feature |  | Count |
| --- | --- | --- | --- |
| 1 | Number of samples (GISAID) |  | 200865 |
| 2 | Total mutations |  | 2583007 |
| 3 | Nonsynonymous mutations |  | 1327548 (51.39 %) |
|  | a | Missense mutation | 657667 (49.54 %) |
|  | b | Stop lost | 15532 (1.17 %) |
|  | c | Stop gain | 8761 (0.66 %) |
|  | d | Deletions/ Insertion | 3053 (0.23 %) |
| 4 | Synonymous mutations |  | 1004453 (38.88 %) |
| 5 | Intergenic mutations |  | 242631 (9.39 %) |
| 6 | Number of unique mutations in the coding region |  | 31641 |
| 7 | Number of unique mutant positions in the coding region |  | 21016 |
| 8 | Number of nonsynonymous mutations per sample (average) |  | 6.6 |
| 9 | Number of synonymous mutations per sample (average) |  | 5 |
| 10 | Number of Intergenic mutations per sample (average) |  | 1.2 |

From the variant dataset generated, we observed 13 hotspot residues across the *SARS-CoV-2* genome that occur at least in 40,000 samples or more in a non-exclusive manner (Figure 1), consistent with literature (15). Among these, the most frequent synonymous mutation c.2772C>T p.F924F occur 186189 times in NSP3 gene (predicted phosphodiesterase) followed by c.14143C>T p.L4715L at 185945 times in RNA-dependent RNA polymerase gene (Table 2). The most frequent nonsynonymous mutations D614G and A222V occur 176436 and 47971 times in the spike glycoprotein S gene, followed by R/G203K/R induced by a tri-nucleotide mutation resulting in a 2-amino acid change, and A220V in the nucleocapsid N gene for 63336 and 48426 times, respectively. Of note, patients infected with the D614G mutation are associated with higher viral loads in the upper respiratory tract than seen with the ancestral strain, but not with altered disease severity (16). However, other reported spike glycoprotein mutations N439K, S477Y, E484K, and N501Y were not found to be significantly abundant in the current data set analyzed. The 13 most recurrent hotspot mutations found comprise of 5 synonymous mutations likely affecting mRNA splicing or selection on codon usage bias, stability and folding translation or co-translational protein folding (17-19), that remains to be explored.

**Table 2:**
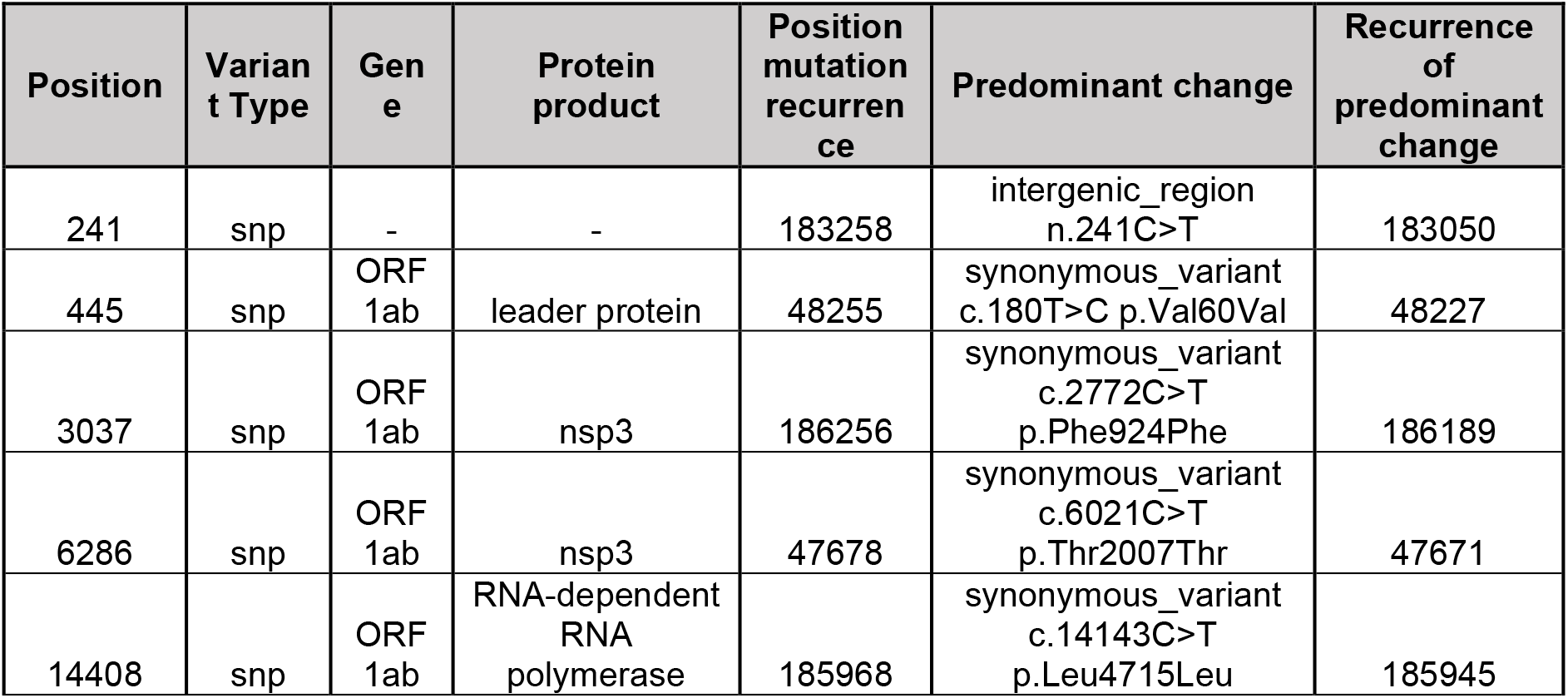
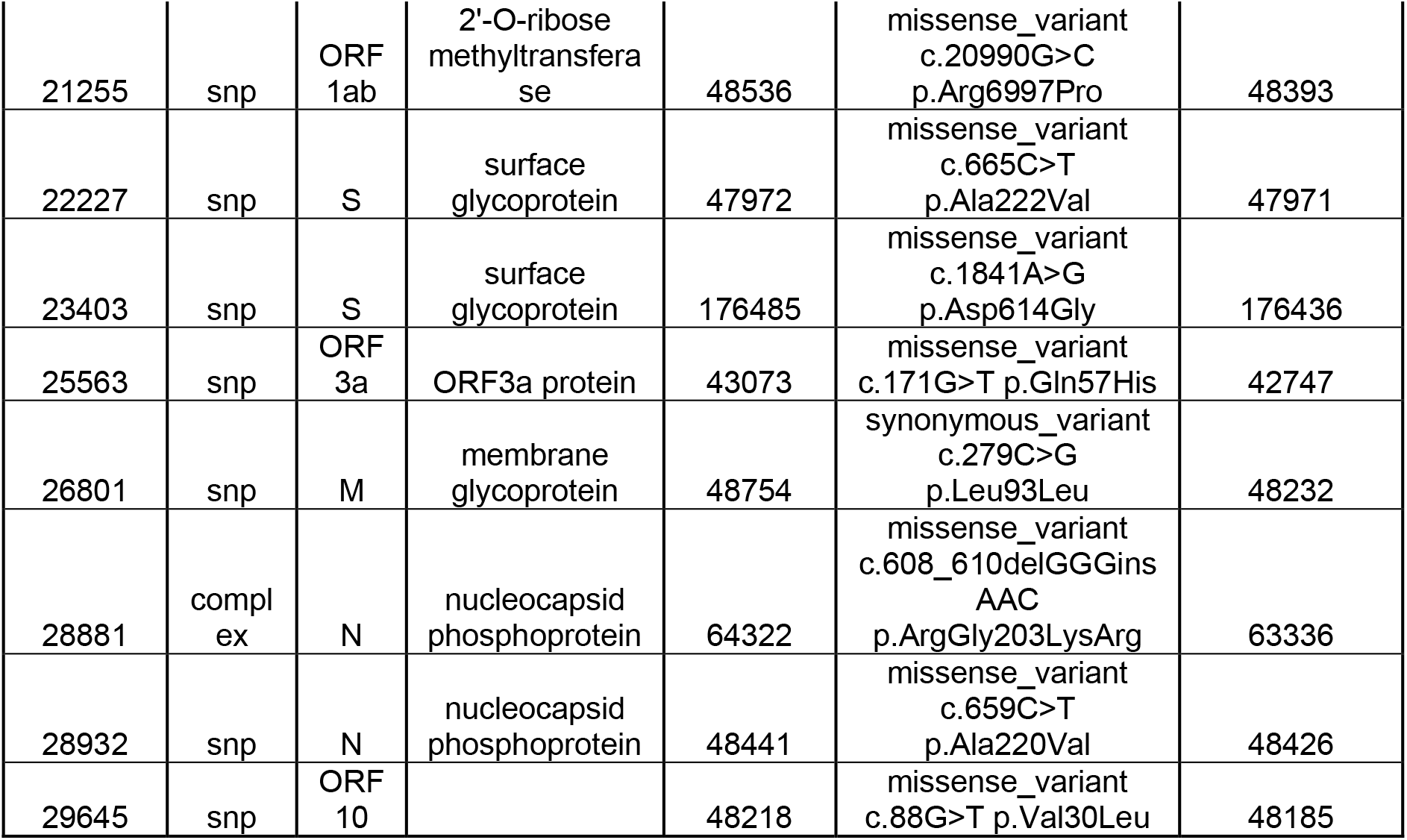
Recurrent hot-spot mutations in the *SARS-CoV-2* genome

| Position | Variant Type | Gene | Protein product | Position mutation recurrence | Predominant change | Recurrence of predominant change |
| --- | --- | --- | --- | --- | --- | --- |
| 241 | snp | - | - | 183258 | intergenic_region<br>n.241C>T | 183050 |
| 445 | snp | ORF1ab | leader protein | 48255 | synonymous_variant<br>c.180T>C p.Val60Val | 48227 |
| 3037 | snp | ORF1ab | nsp3 | 186256 | synonymous_variant<br>c.2772C>T<br>p.Phe924Phe | 186189 |
| 6286 | snp | ORF1ab | nsp3 | 47678 | synonymous_variant<br>c.6021C>T<br>p.Thr2007Thr | 47671 |
| 14408 | snp | ORF1ab | RNA-dependent RNA polymerase | 185968 | synonymous_variant<br>c.14143C>T<br>p.Leu4715Leu | 185945 |
| 21255 | snp | ORF 1ab | 2'-O-ribose methyltransferase | 48536 | missense_variant<br>c.20990G>C<br>p.Arg6997Pro | 48393 |
| 22227 | snp | S | surface glycoprotein | 47972 | missense_variant<br>c.665C>T<br>p.Ala222Val | 47971 |
| 23403 | snp | S | surface glycoprotein | 176485 | missense_variant<br>c.1841A>G<br>p.Asp614Gly | 176436 |
| 25563 | snp | ORF 3a | ORF3a protein | 43073 | missense_variant<br>c.171G>T p.Gln57His | 42747 |
| 26801 | snp | M | membrane glycoprotein | 48754 | synonymous_variant<br>c.279C>G<br>p.Leu93Leu | 48232 |
| 28881 | complex | N | nucleocapsid phosphoprotein | 64322 | missense_variant<br>c.608_610delGGGins<br>AAC<br>p.ArgGly203LysArg | 63336 |
| 28932 | snp | N | nucleocapsid phosphoprotein | 48441 | missense_variant<br>c.659C>T<br>p.Ala220Val | 48426 |
| 29645 | snp | ORF 10 |  | 48218 | missense_variant<br>c.88G>T p.Val30Leu | 48185 |

**Figure 1:**
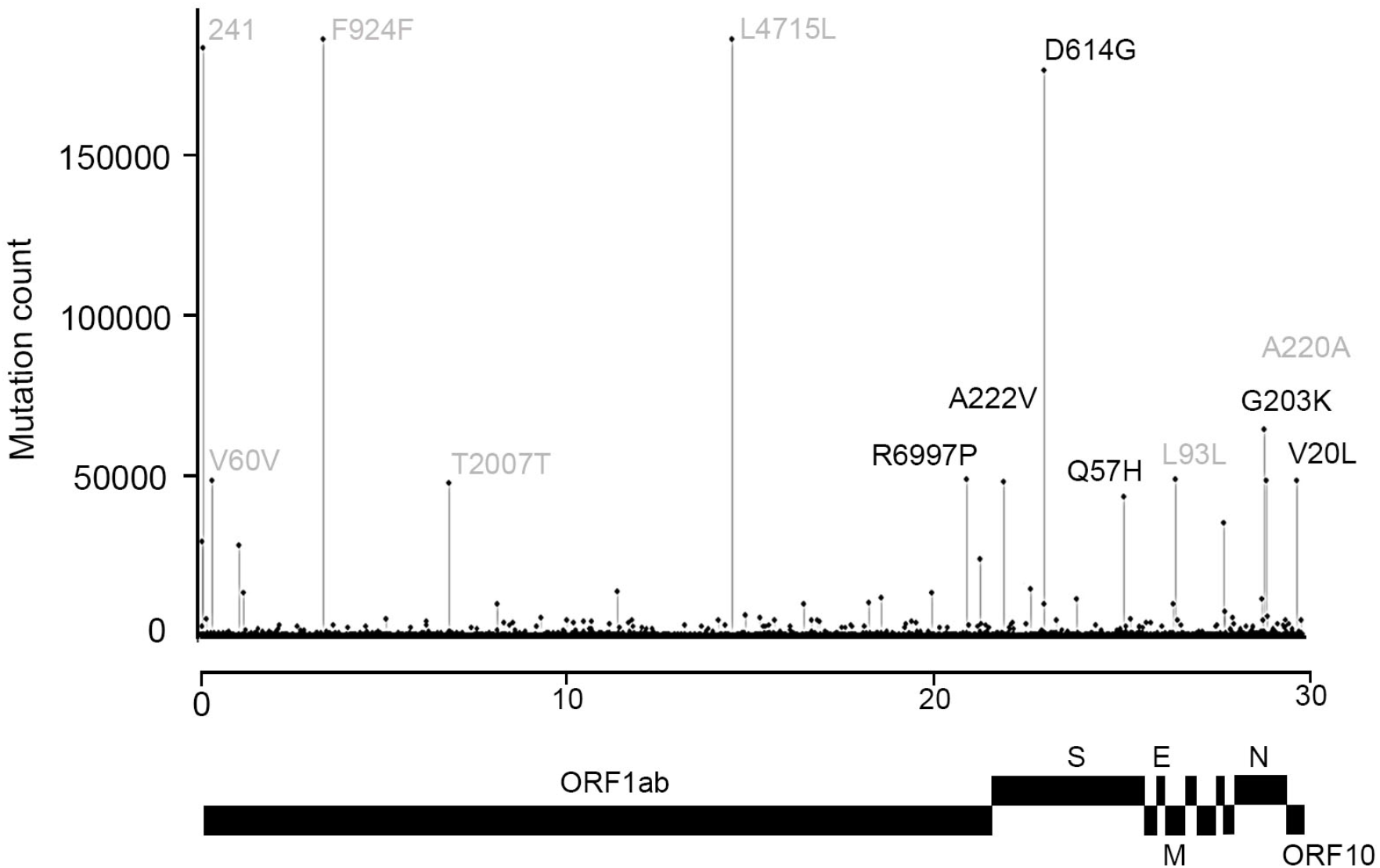
Global distribution of *SARS-CoV-2* genome mutations. The hotspot mutations (recurrence > 40,000 samples) have been labelled with protein change in the plot. The intergenic and synonymous mutations are colored grey. The gene annotation track on the x-axis is not to scale.

Next, we performed a gene-specific analysis from the generated variant data, to estimate frequencies for genes with under-sampled synonymous mutations accounting for the individual gene biases. Our analysis revealed that after normalizing for gene length, the S, N, M, ORF7a, and ORF10 viral genes that comprise of about 21% of the genome accounts for 54.36% of all *SARS-CoV-2* nonsynonymous mutations (Figure 2). Interestingly, S and M genes harbor least proportion of total variable bases across the *SARS-CoV-2* genome indicating a strong positive selection of nonsynonymous mutations in both the genes (Figure 3). The insights of functional relevance of the different amino acid sites mutated though remain to be established.

**Figure 2:**
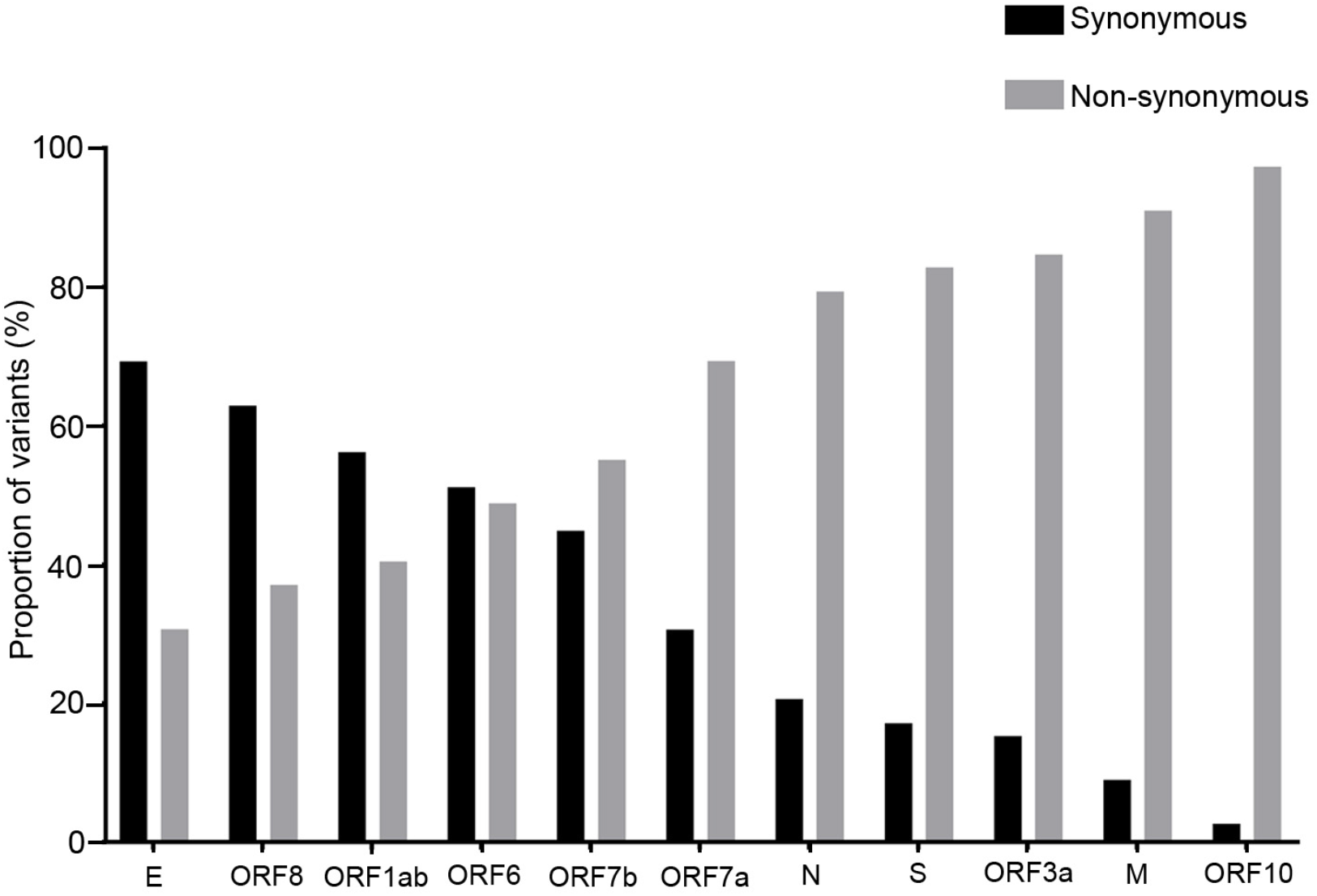
Proportion of synonymous and non-synonymous mutations across all the *SARS-CoV-2* genes

**Figure 3:**
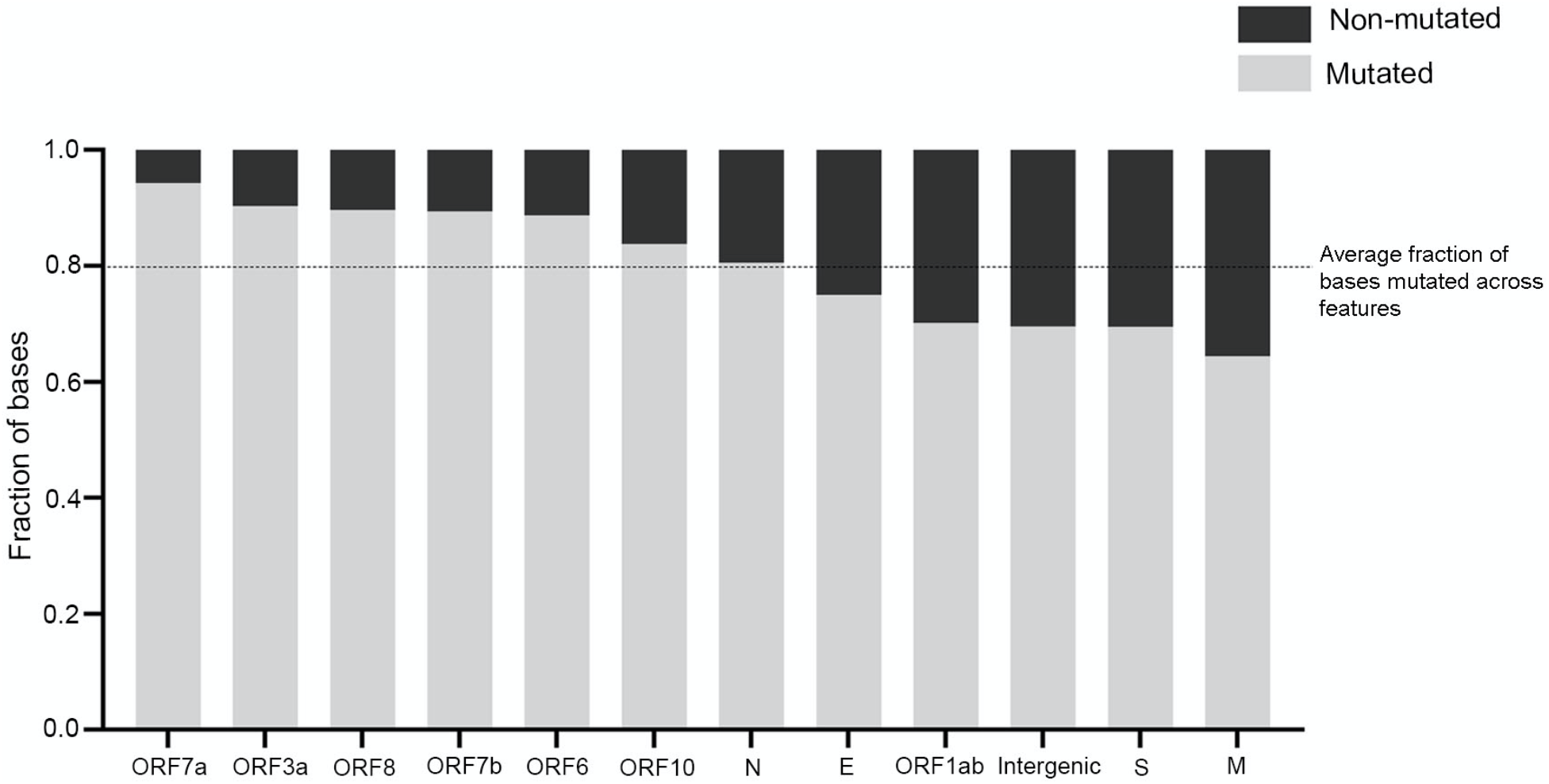
Proportion of mutated/ non-mutated bases across the *SARS-CoV-2* gene features. The dotted line indicates average fraction of mutated residues per feature (∼ 0.8).

We also analyzed for variants in the newer *SARS-CoV-2* virus lineage B1.1.7 (clade 20I/501.V1) emerging in the UK (20), B.1.351 (clade 20H/501Y.V2) in South Africa (21), and P.1 (clade 20J/501Y.V3) in Brazil (12), that were found to harbor a total of 32, 25 and 25 median mutations across 13, 82 and 13 samples, respectively, for each lineage (Table 3 and Supplementary Table 1). A comparative account of variants predominant in the three newer lineages originating from distinct geographical regions along with those reported from India, comprising of 3,361 samples with a comparable frequency of nonsynonymous mutations (48.75%) and synonymous mutations (41.45%) (Table 4), revealed four core common hotspot mutations including D514G mutation in the spike protein and several lineage restricted unique mutations for each strain (Figure 4A). Among the three emergent strains, N501Y was found as the root mutation, while the South African and Brazil strain appear to acquire additional lineages specific E484K mutation within spike protein. Taken together, this suggests a repetitive convergent and adaptive evolution adopted by the distinct lineages (Figure 4B) that tends to pose a reasonable threat towards the emergence of newer regional variant strains with continued persistence of the pandemic. Of note, no significant incidence of N501Y or E484K spike protein mutations, predominant in the emerging strains, occur in the Indian samples. Of earlier reported spike protein mutations, S477N mutation was found in 2 of 3,361 Indian samples, although no incidence of N439K was found. Whether these mutations that show enhanced binding affinity to human and murine ACE2 receptor (22) could account for the exponential transmission rate observed among the newer emergent *SARS-CoV-2* virus lineages in the UK, South Africa and Brazil, but not in India, remains to be established.

**Table 3:**
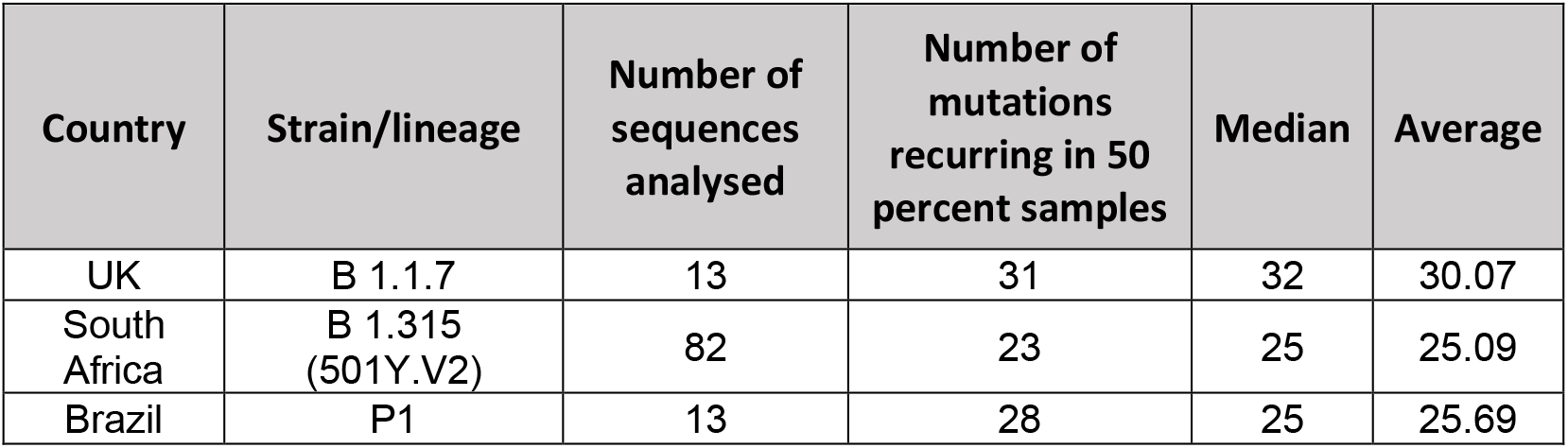
Variant analysis statistics of the emerging SARS-CoV-2 lineages, B 1.1.7 (UK), B 1.315 (South Africa) and P1 (Brazil)

**Table 4:**
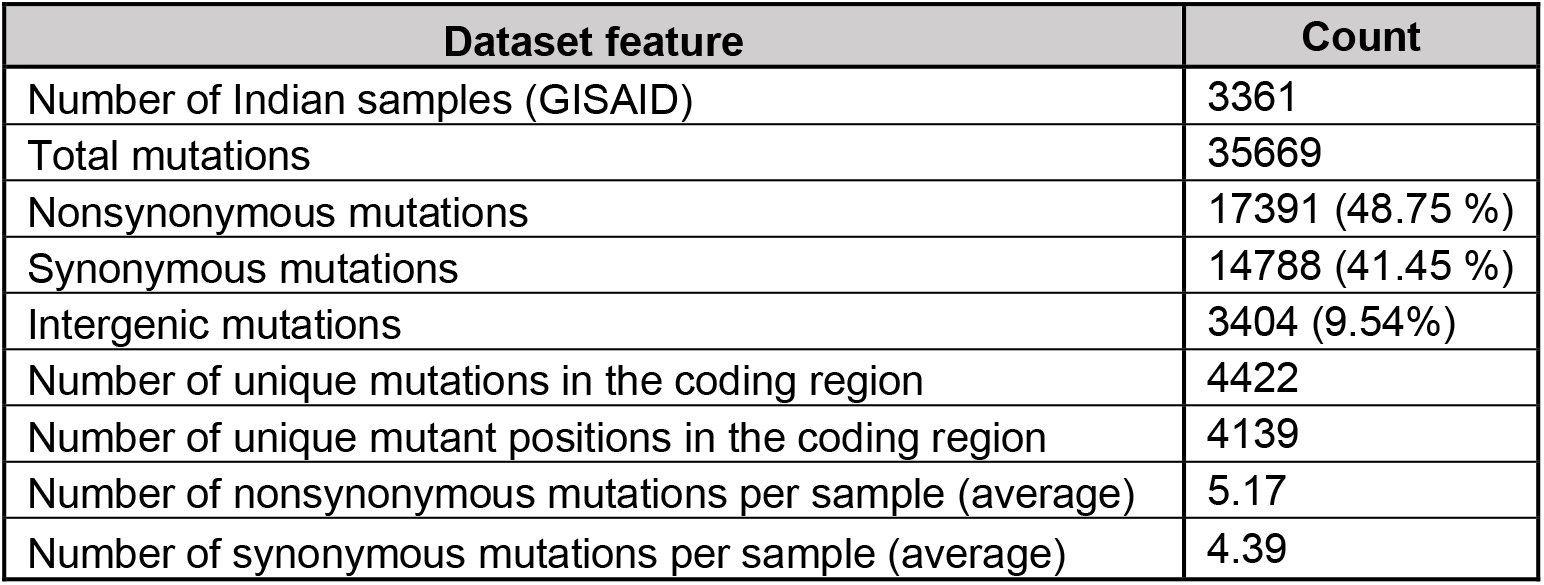
Distribution of type of *SARS-CoV-2* genome variants in Inian samples submitted in GISAID database

**Figure 4:**
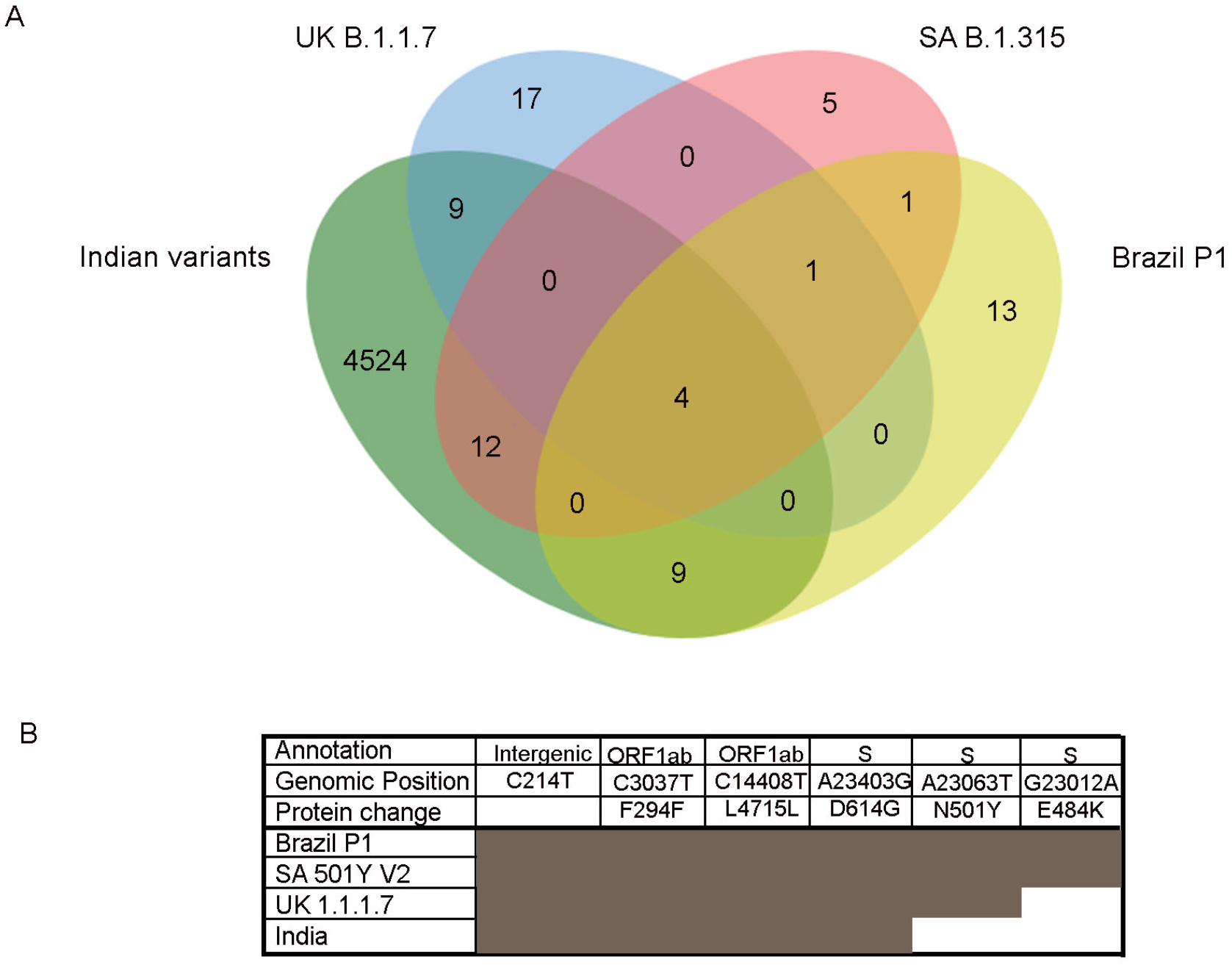
Overlap of variants recurring among the emerging strains (B 1.1.7, B 1.135 and P1) and Indian samples. A. Variants recurring in at least 50 percent of analyzed samples are overlapped with variants in Indian samples. B. Variants common across all the strains, including Indian samples and private clade defining variants in the S protein across the emerging SARS-CoV-2 strains

Upon performing the clade analysis on the downloaded GISAID sequence data, we observed the predominance of 20E (EU1) (26.2%), 20B (26.13%) and 20A (20.27%) clades in the sample set (Figure 5). The variant and clade information generated from the analysis formed the core database for the *SARS-CoV-2* module of IPD 2.0. We tested the clade assessment function of IPD 2.0 using a simulated sequencing dataset generated from representative genomes from different clades (as described in Material and methods). We found that IPD 2.0 could assign the clades with high accuracy.

**Figure 5:**
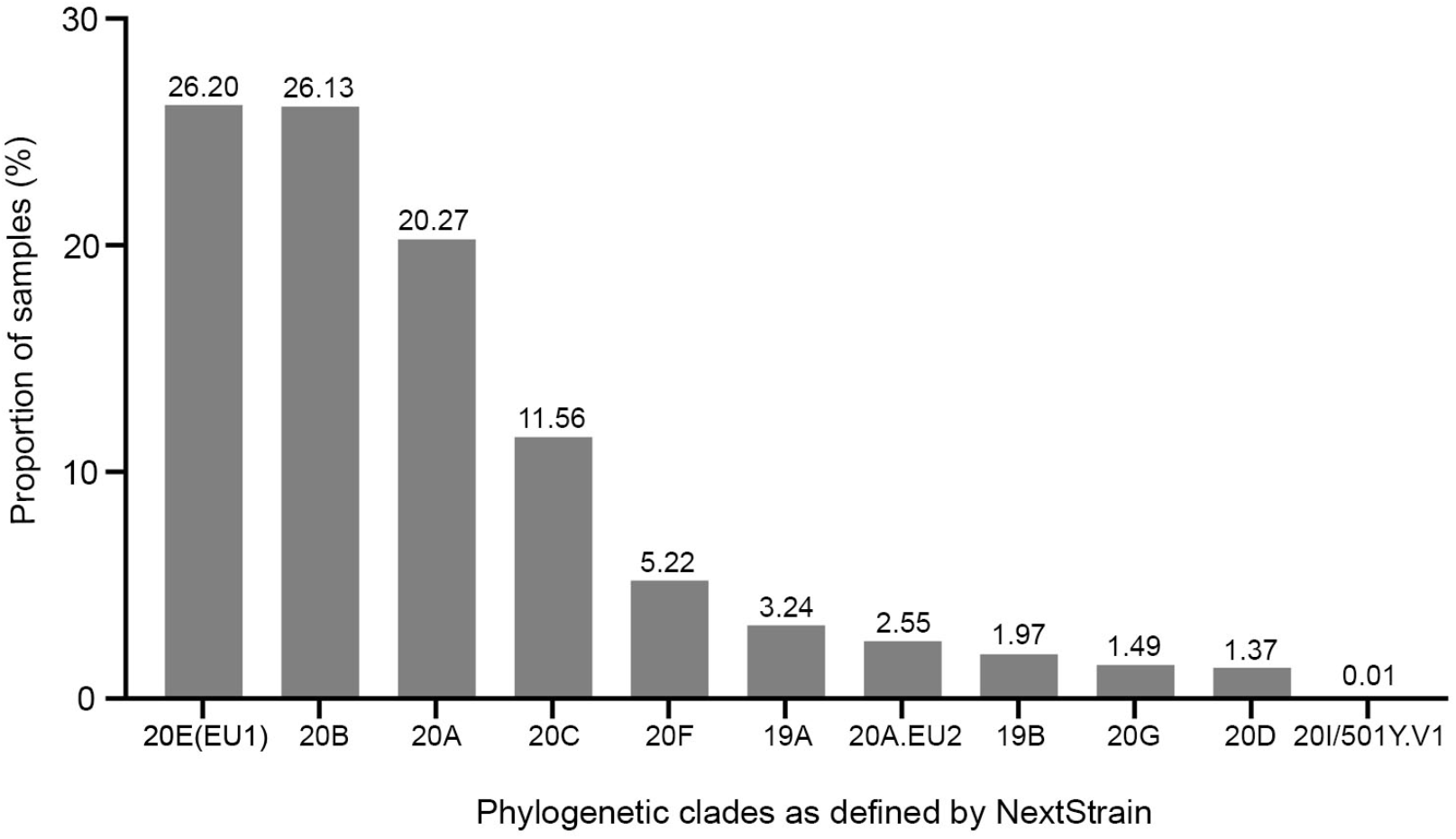
Proportion of samples representing clades in the GISAID samples (n=200865)

In summary, we have developed a pipeline for quantification and phylogenetic assessment of *SARS-CoV-2* genome, which incorporates updated variant and clade assessment module. This makes IPD 2.0 a pertinent tool for analysis of diverse *SARS-CoV-2* sequence datasets and facilitate genomic surveillance to identify variants involved in breakthrough infections.

## Availability and implementation

IPD 2.0 is freely available from http://www.actrec.gov.in/pi-webpages/AmitDutt/IPD/IPD.html and the web-application is available at http://ipd.actrec.gov.in/ipdweb/

