## Supplementary methods for "Evolving Insights from *SARS-CoV-2* Genome from 200K COVID-19 Patients"

### Generation of simulated test data for clade assessment of IPD 2.0

To evaluate accuracy of the modified SARS-CoV-2 module, clade assessment of IPD 2.0, test dataset using the sequences downloaded from GISAID. Clade assessment of these sequences was performed using NextClade analysis of the NextStrain package. The test data set consisted of total 16 simulated samples (8 with introduced background mutation and 8 without background mutations inserted), generated using the neatgenreads tool. The samples generated were of 10x coverage and having read length of 101 bp. In the Random mutation insertion = 0 and 0.2

We used the following SARS-CoV-2 reference genomes from different clades (determined by Nextclade analysis.)

1. hCoV-19/Japan/NA-20-05-1/2020|EPI_ISL_410531|2020-01-25 (clade 19A)
2. hCoV-19/Singapore/4/2020|EPI_ISL_410535|2020-02-03 (clade 19B)
3. hCoV-19/Germany/SL-SU-10429159/2020|EPI_ISL_707956|2020-03-24 (clade 20A.EU2)
4. hCoV-19/England/NORW-E89D6/2020|EPI_ISL_448260|2020 (clade 20A)
5. hCoV-19/England/NORW-E8A1F/2020|EPI_ISL_448264|2020 (clade 20B)
6. hCoV-19/Netherlands/GE-EMC-250/2020|EPI_ISL_523229|2020 (clade 20C)
7. hCoV-19/Israel/CVL-n-5939/2020|EPI_ISL_474965|2020-03-15 (clade 20D)
8. hCoV-19/Gibraltar/204641742/2020|EPI_ISL_637212|2020 (clade 20E.EU1)
