## Supplementary Table 1 for "Evolving Insights from *SARS-CoV-2* Genome from 200K COVID-19 Patients"

**Supplementary Table S1**: Variants identified in at least 50 percent of the strain specific sequences analysed from P.1, B 1.315 and B 1.1.7 strain

| P1 Brazil strain variants | B 1.315 South Africa strain variants | B 1.1.7 UK strain variants |
|---|---|---|
| A5648C | A10323G | A23063T |
| A6613G | A23063T | A23403G |
| AGTAGGG28877TCTAAAC | A23403G | A28111G |
| C14408T | C1059T | ATACATG21764A |
| C241T | C14408T | C14408T |
| C28512G | C23664T | C23604A |
| C3037T | C241T | C23709T |
| G17259T | C25904T | C27972T |
| G25088T | C26456T | C3037T |
| G28262GAACA | C28887T | G24914C |
| T26149C | G174T | G28048T |
| T733C | G22813T | GGG28881AAC |
| A6319G | G23012A | C241T |
| C12778T | G25563T | C28977T |
| C13860T | G5230T | C5388A |
| C21614T | A21801C | C5986T |
| C21621A | C3037T | GAT28280CTA |
| C21638T | C14925T | TA28270T |
| C24642T | C28253T | C14676T |
| A23403G | C8964T | C15279T |
| C23525T | G11230T | C23271A |
| C2749T | C8660T | C3267T |
| G21974T | T27504C | C913T |
| G28167A | | GTCTGGTTTT11287G |
| G22132T | | T16176C |
| A22812C | | T24506G |
| A23063T | | T6954C |
| G23012A | | TTTA21990T |
